# Morphogen dynamics control patterning in a stem cell model of the human embryo

**DOI:** 10.1101/202366

**Authors:** Idse Heemskerk, Kari Burt, Matthew Miller, Sapna Chhabra, M. Cecilia Guerra, Aryeh Warmflash

**Affiliations:** Department of Biosciences, Rice University, Houston TX 77005.; Systems, Synethetic and Physical Biology Program, Rice University, Houston TX 77005.; Department of Bioengineering, Rice University, Houston TX 77005.

**Author notes:** Correspondence to AW.

## Abstract

During embryonic development, diffusible signaling molecules called morphogens are thought to determine cell fates in a concentration-dependent manner^1–4^, and protocols for directed stem cell differentiation are based on this picture^5–8^. However, in the mammalian embryo, morphogen concentrations change rapidly compared to the time for making cell fate decisions^9–12^. It is unknown how changing ligand levels are interpreted, and whether the precise timecourse of ligand exposure plays a role in cell fate decisions. Nodal and BMP4 are morphogens crucial for gastrulation in vertebrates^13^. Each pathway has distinct receptor complexes that phosphorylate specific signal transducers, known as receptor-Smads, which then complex with the shared cofactor Smad4 to activate target genes^14^. Here we show in human embryonic stem cells (hESCs) that the response to BMP4 signaling indeed is determined by the ligand concentration, but that unexpectedly, the expression of many mesodermal targets of Activin/Nodal depends on rate of concentration increase. In addition, we use live imaging of hESCs with GFP integrated at the endogenous *SMAD4* locus to show that a stem cell model for the human embryo^15^ generates a wave of Nodal signaling. Cells experience rapidly increasing Nodal specifically in the region of mesendoderm differentiation. We also demonstrate that pulsatile stimulation with Activin induces repeated strong signaling and enhances mesoderm differentiation. Our results break with the paradigm of concentration-dependent differentiation and demonstrate an important role for morphogen dynamics in the cell fate decisions associated with mammalian gastrulation. They suggest a highly dynamic picture of embryonic patterning where some cell fates depend on rapid concentration increase rather than absolute levels, and point to ligand dynamics as a new dimension to optimize protocols for directed stem cell differentiation.

---

In the early mammalian embryo, BMP4 defines the dorsal-ventral axis while Nodal initially maintains the pluripotent epiblast and is subsequently required for mesoderm and endoderm differentiation. Expression patterns of these morphogens change significantly through gastrulation, which also involves large scale cell movements, and therefore individual cells experience rapidly changing morphogen levels during differentiation. To investigate how sudden increases in morphogen levels are interpreted by hESCs we focused on SMAD4, a shared component of the BMP and Nodal pathways (Fig. 1a) that translocates to the nucleus when the pathways are active. We performed live imaging of cells with GFP:SMAD4 in the endogenous locus^16^, and quantified signaling strength as the ratio of nuclear and cytoplasmic SMAD4 intensity (Extended Data Fig. 1a). We found that a sudden increase in BMP4 leads to sustained Smad4 signaling (Fig. 1b,c). In contrast, the response to addition of the Nodal substitute Activin is strongly adaptive, i.e. transient, and returns to a signaling baseline of around 20% of the response peak (Fig 1b,d), similar to the previously observed response to TGFβ in C2C12 cells^17,18^. The amplitude and baseline, but not the timescale of adaptation depended on the Activin dose (Extended data Fig 1b), while for BMP4 the dose affected the duration of signaling in a way consistent with ligand depletion at low doses (Extended data Fig 1c). Immunofluorescence staining for receptor-Smads revealed that SMAD1 activation follows the SMAD4 response to BMP4, while in response to Activin, SMAD2 nuclear localization adapts less than SMAD4 to about 60% of peak response (Extended data Fig. 1f-j).

**Figure 1.**
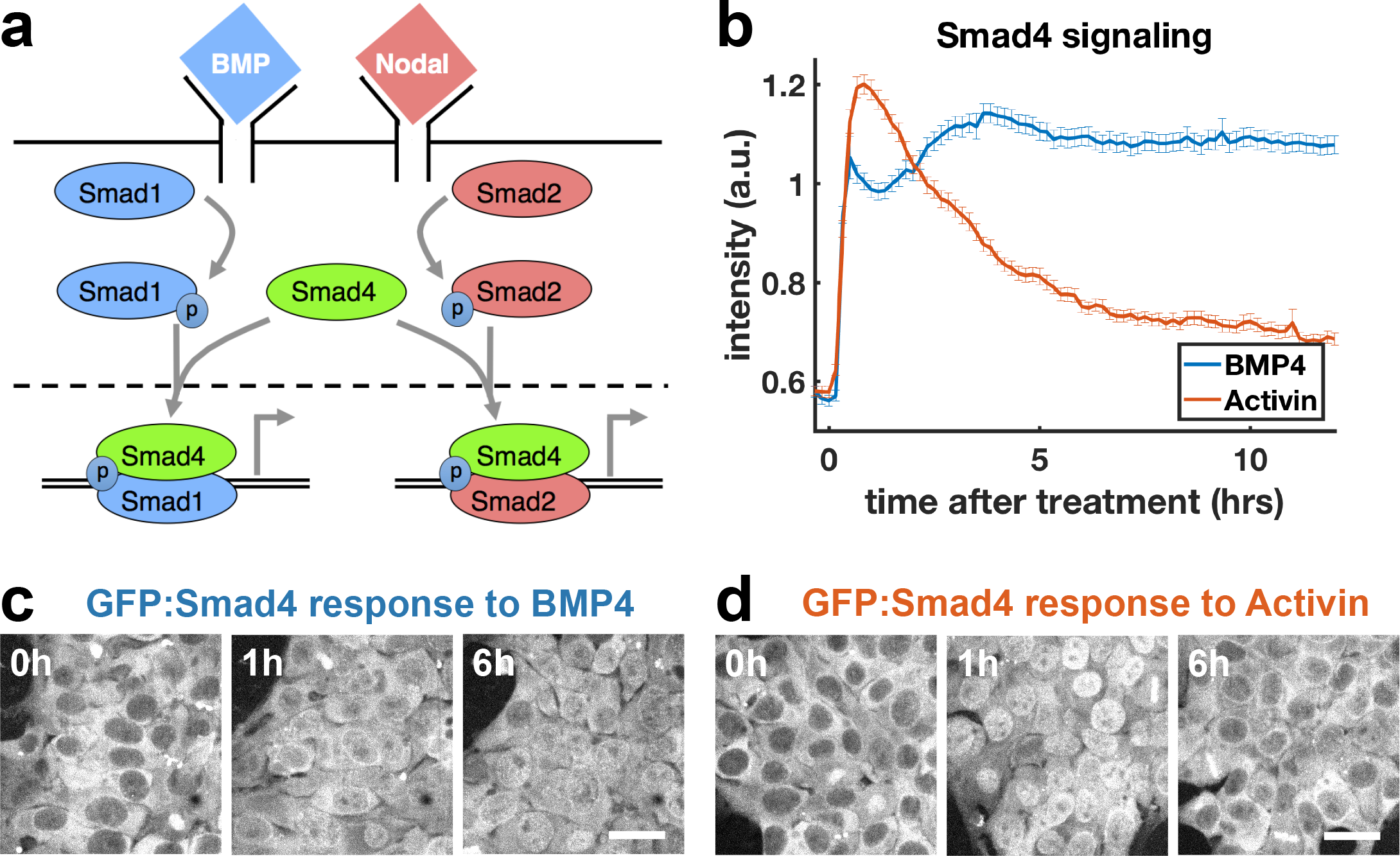
SMAD4 signaling response of hESCs to BMP4 is sustained while that to Activin is adaptive. **a,** BMP and Nodal pathways share the signal transducer Smad4. **b,** GFP:SMAD4 average nuclear intensity after treatment with BMP4 (blue) or Activin (red). Error bars represent standard error. (Ncells~700). Distributions shown in Extended data Fig.1d-e. **c,d,** hESCs expressing GFP:SMAD4 at 0, 1 and 6 hours after treatment with BMP4 **(c)** or Activin **(d)**. Scalebar 50μm.

Next, we evaluated transcriptional dynamics downstream of BMP4 and Activin using qPCR, which showed that BMP targets are stably induced (Fig. 2a), while differentiation targets of Nodal show adaptive transcription on a timescale consistent with SMAD4 signaling (Fig. 2b). Moreover, shared targets of the pathways were found to be transcribed adaptively in response to Activin and stably in response to BMP4 (Fig. 2c, Extended data Fig. 2a). In contrast, the transcription of *NODAL*, *WNT3*, and their inhibitors LEFTY1 and CER1 were sustained (Fig. 2d). The sustained transcription of these targets may reflect a requirement to activate the positive feedbacks between the Nodal and Wnt pathways, which are known to be involved in establishing the primitive streak, the region of the mammalian embryo where mesoderm and endoderm form^19^. This suggests a picture where stable transcription of the ligands and inhibitors allows for the establishment of signaling patterns in the embryo while cells receiving these signals to differentiate interpret them according to their dynamics. Several other genes not related to mesendoderm differentiation were also found to be stably induced by Activin (Extended data Fig. 2b-d). Molecularly, the two classes of transcriptional dynamics may reflect differential requirements for SMAD4 signaling levels with lower levels required to maintain the targets with sustained dynamics, or SMAD4 independent transcription that depends only on SMAD2 activation, which is more sustained (Extended data Fig 1f-j). Because Nodal maintains epiblast pluripotency in combination with FGF^20,21^ but induces primitive streak fates when Wnt is present, we then asked whether target genes respond adaptively to Nodal signaling during differentiation. Initial transcriptional response was found to be qualitatively similar with or without Wnt activation, although shared targets of Wnt and Activin such as the mesendodermal markers *EOMESODERMIN (EOMES)* and *BRACHYURY (BRA)* showed stronger response and could be stably reactivated following adaptation on longer time scales (Fig. 2e, Extended data Fig. 2e,f).

**Figure 2.**
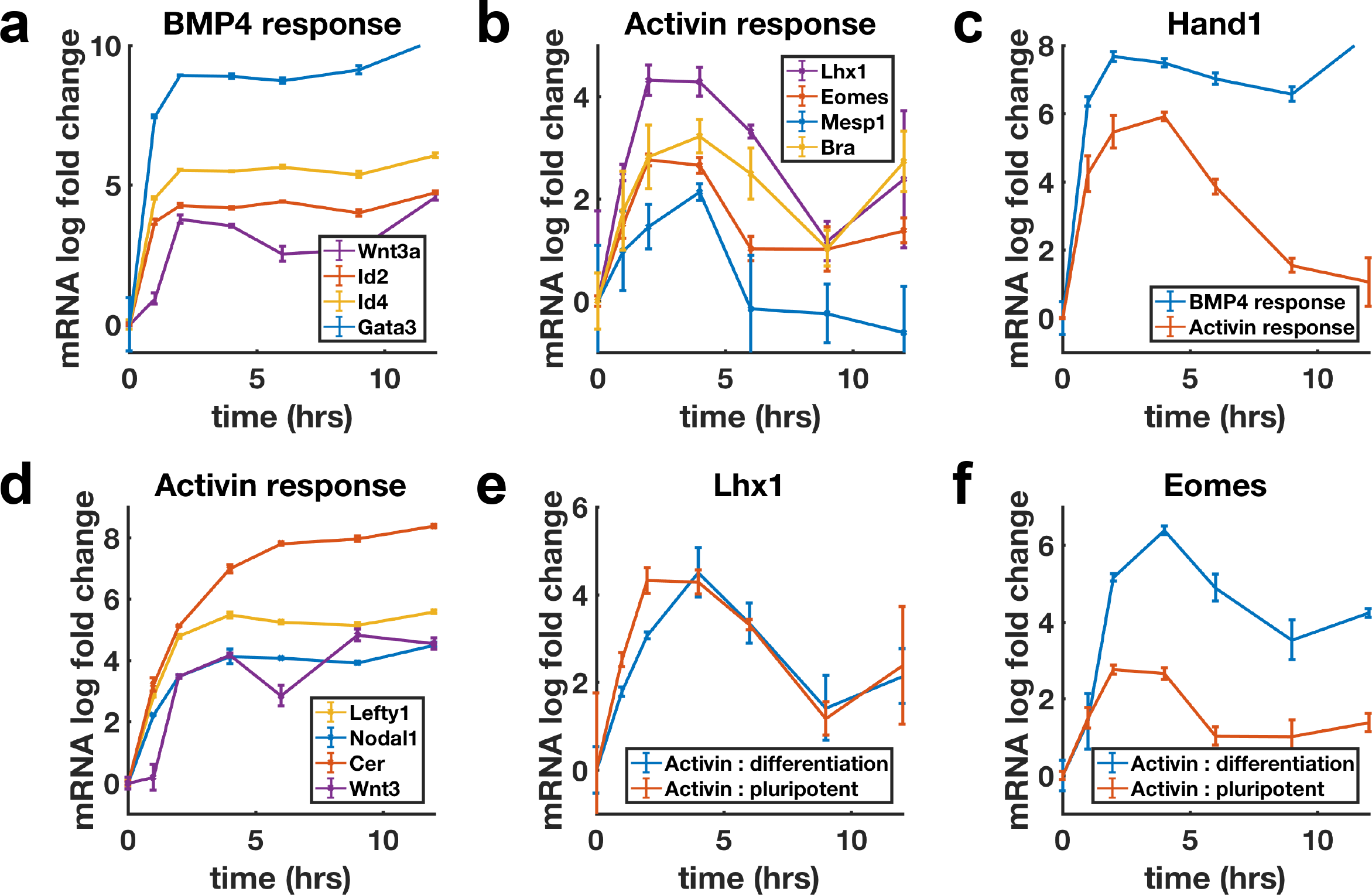
Transcription of BMP targets and Nodal differentiation targets reflects Smad4 dynamics, while other Nodal targets show sustained transcription. **a, b,** qPCR measurements of transcriptional response to BMP4 treatment **(a)** and of differentiation targets to Activin **(b),** y-axes show relative C_T values. **c,** Transcription of shared Activin/BMP4 target *HAND1* after BMP4 (blue) or Activin (red) treatment. **d,** Non-adaptive response to Activin of morphogens and inhibitors involved in initiating the primitive streak. **e,** Transcriptional response to Activin under pluripotency maintaining conditions (red) and mesendoderm differentiation conditions (blue) of Activin target *LHX1* **(e)** and joint Activin/Wnt target *EOMES* **(f)**.

Sustained response to BMP4 suggests sensing of ligand concentration. In contrast, adaptive response to Activin with dose dependent amplitude suggests sensing of ligand rate of increase. We tested this by slowly raising Activin and BMP4 concentrations (concentration ramp) and comparing the response to these ramps with the response to a single step to the same final dose (Fig. 3a,d). If cells are primarily sensitive to ligand doses, the step and ramp should eventually approach the same final activity, while if cells are sensitive to the rate of ligand increase, the response to the ramp should be reduced. As expected, BMP4 signaling responses to the ramp and step approached each other as ramp concentration increased, while Activin signaling was dramatically reduced in the case of the ramp (Fig 3b,e). Moreover, transcriptional dynamics of the shared target *HAND1* matched the signaling pattern and showed dramatically reduced transcription in response to the Activin ramp. Similar results were obtained for other adaptive Activin targets, in contrast to non-adaptive Activin targets which as expected also showed sustained transcription in response to the ramp (Extended data Fig. 3).

**Figure 3.**
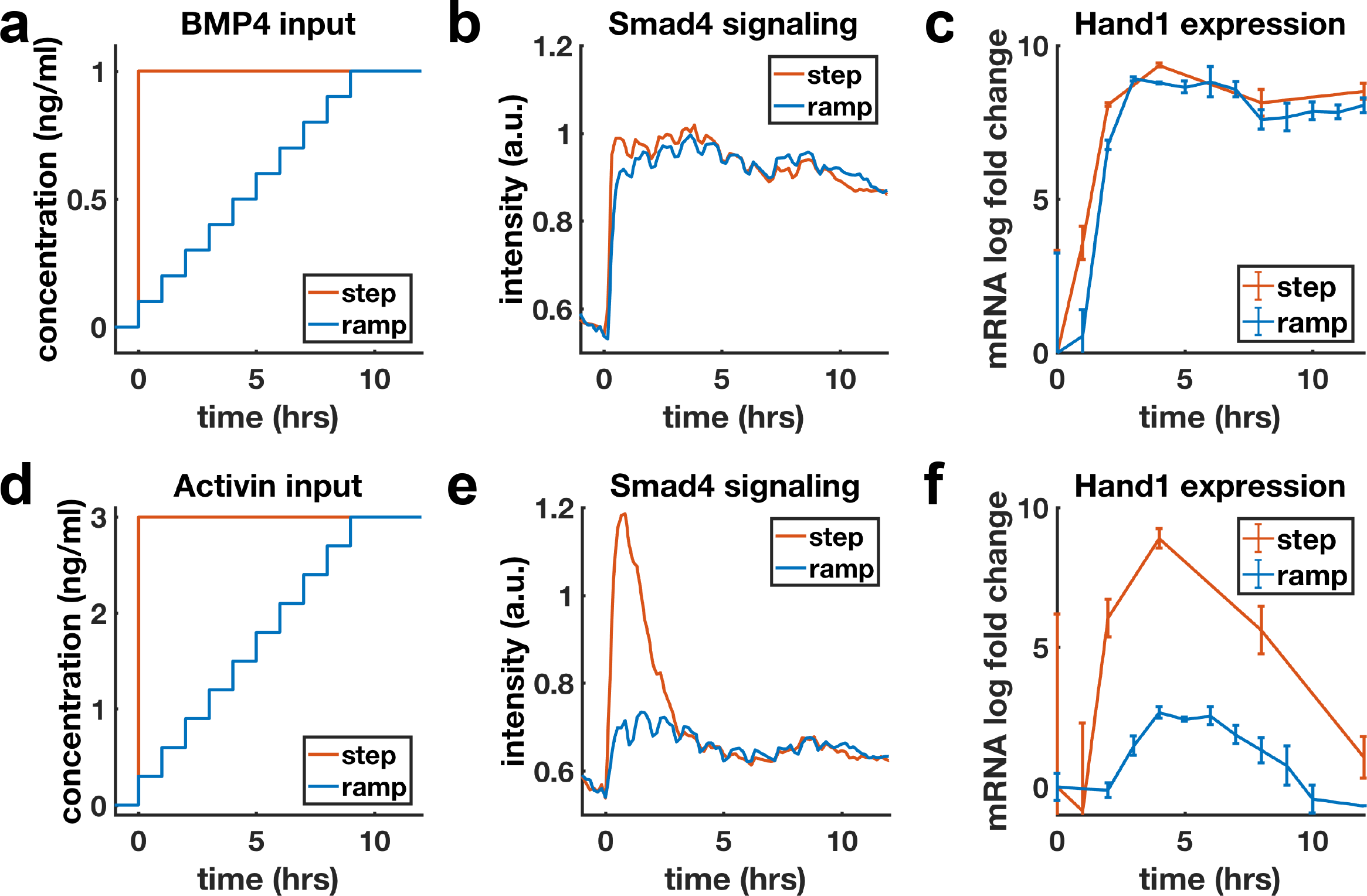
BMP4 response reflects concentration, but Activin response reflects rate of concentration increase. **a, d,** Ligand concentration over time for slow ramp (blue) or sudden step (red) in BMP4 **(a)** or Activin **(d)**. **b, e,** SMAD4 signaling response to BMP4 **(b)** or Activin **(e)**. **c, f,** Transcriptional response of *HAND1* to concentration ramp versus step for BMP4 **(c)** or Activin **(f)**.

Morphogens control cell fate and the dependence of transcription of mesodermal and endodermal genes on Activin rate of increase suggests rapid Activin increase may boost differentiation to these fates. We hypothesized that exposing cells to repeated rapid increases by pulsing the level of Activin could enhance this effect. To rigorously test this hypothesis, we compared pulses that switch between high and low doses of Activin with performing media changes at the same times but maintaining a high Activin dose (“dummy pulses”) (Fig. 4a). Each pulse of Activin elicited a strong response in the translocation of SMAD4 to the cell nucleus while no such responses were seen in response to the dummy pulses (Fig 4b). Differentiation to mesoderm and endoderm as marked by Bra expression was also enhanced under pulsed conditions (Fig. 4c,d), despite the reduced integrated ligand exposure. Continuous exposure to lower doses of Activin and treating cells with Activin for the same total time as the three pulses but in a single pulse showed that the effect is specific to pulsed Activin and not a consequence of simply reducing the integrated Activin exposure (Extended data 4).

**Figure 4.**
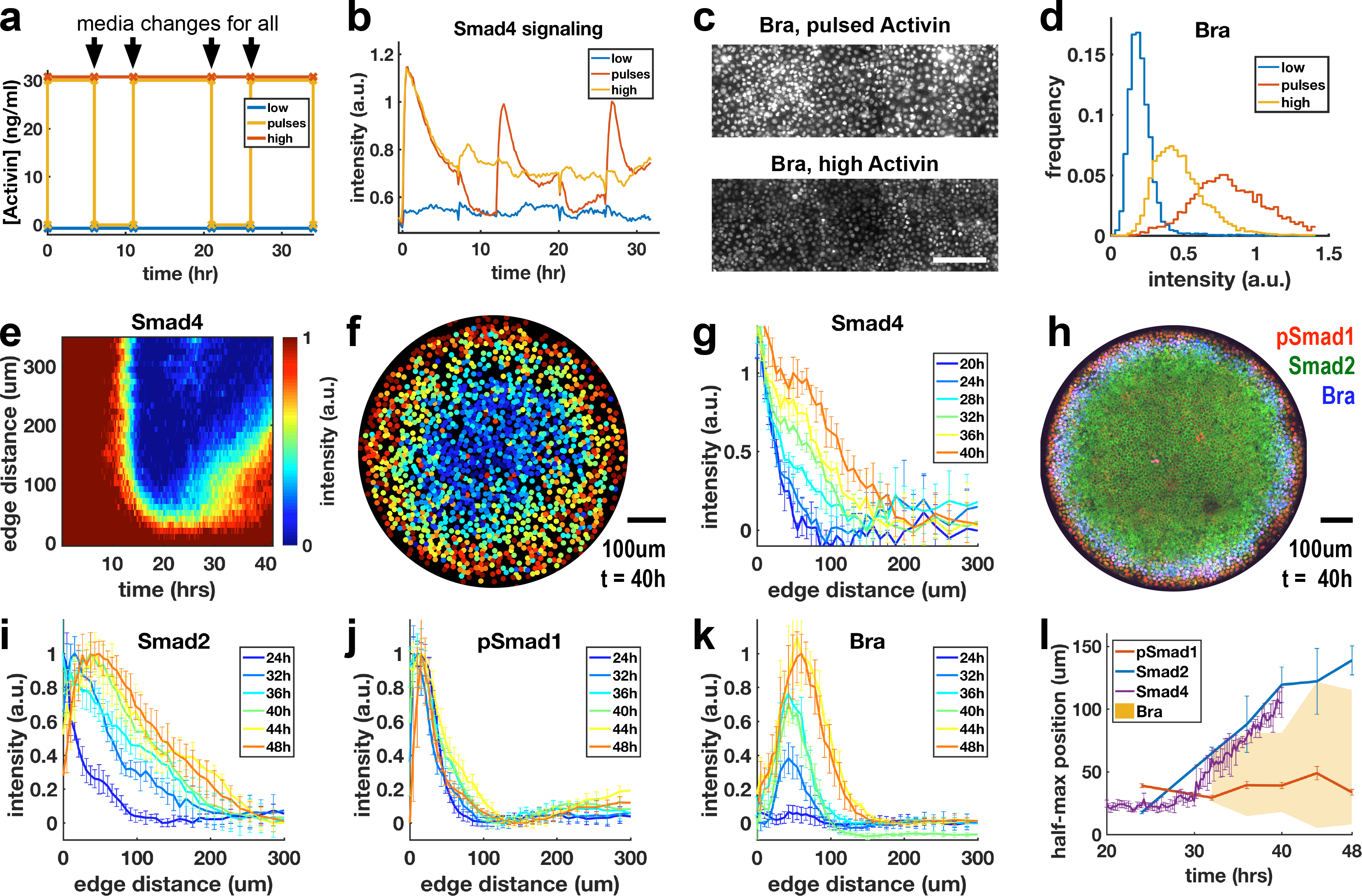
Rapid increases in Activin/Nodal enhance mesoderm differentiation and occur endogenously during self-organized patterning. **a,** Schematic of pulse experiment, graph shows ligand concentration, controls receive media changes at the same time as the pulsed well. **b,** SMAD4 signaling profile in response to Activin pulses (red), high Activin (yellow), and no Activin (blue). **c,** Immunofluorescence staining for BRA after high constant Activin or pulsed Activin. Scalebar 100μm. **d,** Distribution of BRA expression per cell (N_cells_ per condition ~6×10^3^) in images containing **(c)**. **e,** Average radial profile of SMAD4 signaling over time (kymograph) in micropatterned colonies after BMP4 treatment (N=4 colonies). **f,** SMAD4 signaling in single colony at 40h. **g,** Radial SMAD4 signaling profiles at discrete times from 20h to 40h. **h,** Immunofluorescence staining 40h after BMP4 treatment for pSMAD1, SMAD2/3 and BRA. **i,j,k** Normalized radial profiles of SMAD2/3 **(i),** pSMAD1 **(j)** and BRA **(k)** averaged over N>5 colonies per time. **l,** Half-maximum versus time for SMAD2/3 (blue), SMAD4 (purple) and pSMAD1 (red) over time and BRA expression domain defined by a threshold of at least 20% of maximal expression (yellow). In all figures error bars represent standard deviations taken over different colonies.

To test whether rapid concentration increases are relevant to endogenous Nodal during embryonic patterning, we turned to micropatterned colonies of hESCs treated with BMP4. These colonies differentiate in a spatial pattern with reproducible rings of extraembryonic cells and all three germ layers, and represent a model for the patterning associated with gastrulation in the human embryo^15,22^. For the dynamics described here to be relevant, Nodal signaling should evolve rapidly compared to the timescale for adaptation. We used live imaging to observe SMAD4 signaling during the 42 hours in which these patterns form (Fig 4e-g). The cells initially respond uniformly to the BMP4 treatment and this response is restricted to the colony edge by approximately 12 hours (Fig. 4e,f), after which a wave of increased nuclear SMAD4 spreads inward again from the edge (Fig. 4e,g). Because SMAD4 convolves the BMP and Nodal responses, we then looked at the pathway specific SMAD1 and SMAD2 (Fig 4 h-j). These showed that the BMP-specific active SMAD1 signaling remains restricted to the edge (Fig 4 j), while the Nodal transducer SMAD2 activity spreads rapidly inward from the colony edge between 24 and 36h (Fig 4i). Thus, both the SMAD2 and SMAD4 signal transducers reveal a rapidly evolving Nodal distribution with a wavefront that moves through the colony at approximately 1 cell diameter (10μm) per hour. Importantly, this wave of signaling activates Bra expression in its wake and the Bra-positive cells are restricted to the region of the colony with rapid increases in Nodal signaling (Fig. 4k,l). This indicates that rapid increases in endogenous Nodal signaling are associated with mesoderm differentiation in this model of human gastrulation.

Our work shows that morphogens in the mammalian embryo do not act in a purely concentration dependent matter and has revealed an important role for Activin/Nodal dynamics in specifying cell fate. Adaptive signaling could serve to restrict the response to a narrow competence window. Another possible function is the separation of distinct roles for the same morphogen by using distinct dynamics to selectively activate different target genes, effectively expanding the information content of the morphogen gradient^23,24^. It will be important to understand how our results fit with current protocols for directed stem cell differentiation which typically perform a single media change per day. The results presented here will serve as a basis for a dynamic understanding of embryonic patterning and stem cell differentiation.

## Methods

### Cell culture and differentiation protocols

The cell lines used were ESI017 (ESIBIO) and RUES2 GFP:Smad4 RFP:H2B (a gift of Ali Brivanlou, Rockefeller). These cell lines and their maintenance are described in^16^. For all experiments except micropatterning, cells were seeded at a low density of 6×10^4^ cells per cm^2^ and grown with rock-inhibitor Y27672 (10uM; StemCell Technologies). This ensured uniform response to exogenous ligand and minimized the effect of secondary endogenous signaling. Unless otherwise indicated in the figures, experiments in Fig.1–3 were done in mTeSR1 medium (StemCell Technologies) (also referred to as pluripotency conditions), and cells were treated with 50ng/ml Activin A (R&D Systems) or 50ng/ml BMP4 (R&D Systems). Differentiation conditions in Fig.2e are defined as Essential 6 medium (Gibco) + 3uM CHIR99021 (StemCell technologies).

### Imaging and image analysis

Live imaging was done on an Olympus/Andor spinning disk confocal microscope with a 40x, NA 1.25 silicon oil objective. Immunofluorescence data for Fig.4c,d and extended data Fig.4a-e were collected using a 20x, NA 0.75 objective on an Olympus IX83 inverted epifluorescence microscope. Fixed micropatterned colonies for Fig. 4h-k were imaged using an 30x NA 1.05 silicon oil objective on an Olympus FV12 laser scanning confocal microscope. All image analysis used Ilastik^25^ for initial segmentation and custom written Matlab code available on github.com/idse/stemcells for further analysis. Extended Fig.1a shows how Smad4 nuclear to cytoplasmic ratio, our measure for signaling intensity, is defined from the segmentation by subtracting the mean value in the background mask from nuclear and cytoplasmic masks for each cell.

### qPCR

qPCR experiments were done with ESI017 cells using standard methods. Below is a table of primers that were used.

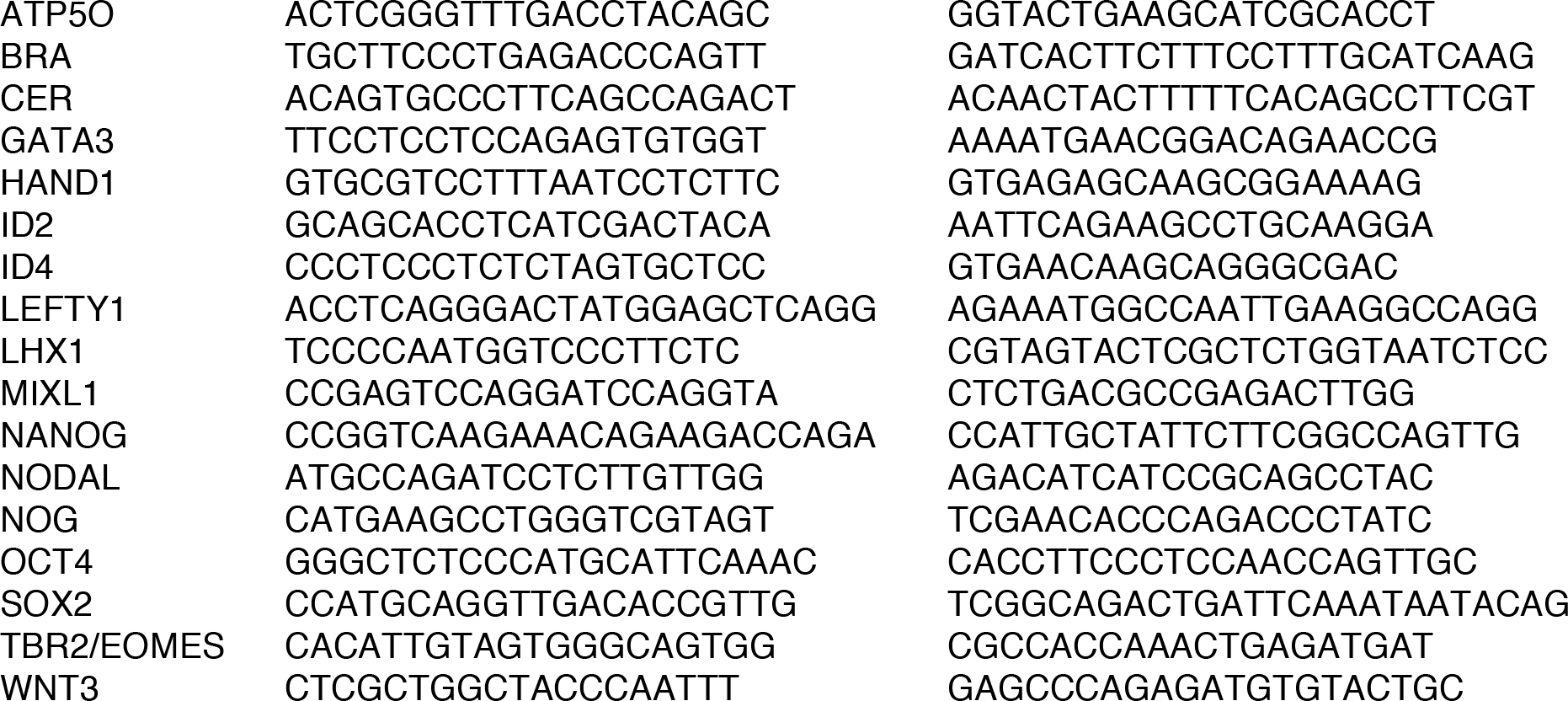

### Pulses

For pulse and dose response experiments in Fig.4 and extended data Fig.4 differentiation was done in Essential 6 medium + 1uM CHIR99021 + 20ng/ml bFGF (Life Technologies) + 10uM Y27672 with 30 ng/ml Activin added as indicated in the figures. Time between pulses was always 5 hours to allow the pathway to relax. Time of individual pulses was chosen for experimental convenience and lengths of pulses shown in Fig.4 are 6, 10, and 8 hours. Controls were subjected to media changes at the same time. During media changes cell were washed 3 times with PBS.

### Micropatterning

For micropatterned colonies we followed the protocol in^26^ using the chemically defined medium mTeSR1. For fixed micropatterns we used the CYTOO Arena A chip and analyzed the 800um colonies, while for live imaging we used a CYTOO 96 well plate RW DS-S-A, which has 700um colonies. For the analysis in Fig.4, radial profiles of SMAD1 and SMAD2 were normalized to have the lowest signaling level be zero and the highest be one in at each time, as minimal and maximal levels were similar at each time and only their spatial distribution was changing. SMAD4 was normalized to have the maximum past position of the half-maximum of SMAD1 be one, as this reflects the peak of Nodal-dependent SMAD4 signaling. Finally, because BRA levels change substantially, BRA at all times was normalized to the maximum of the latest time (48h).

### Immunostaining

Fixing of cells and immunostaining of was done as described in^16^. Below is a table of antibodies that were used.

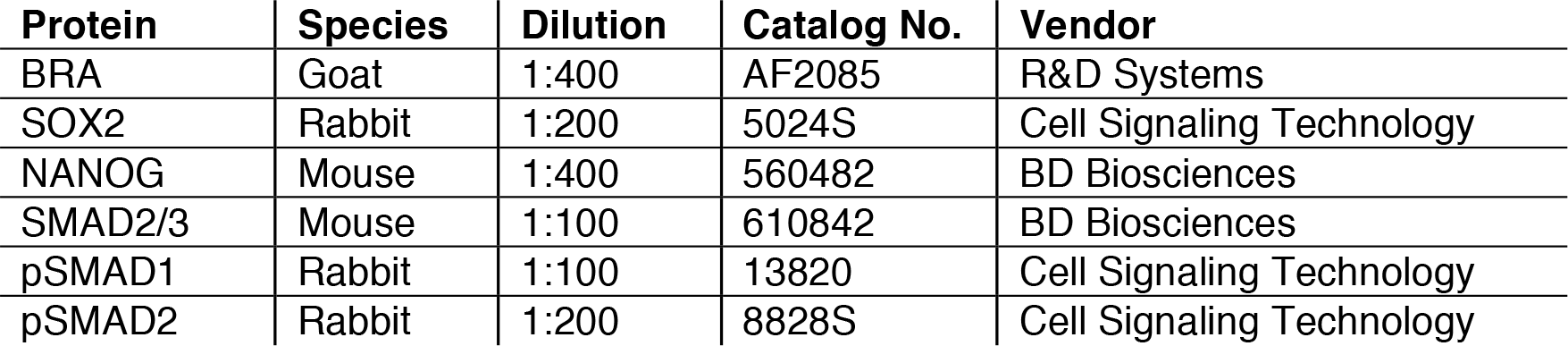

| Protein | Species | Dilution | Catalog No. | Vendor |
| --- | --- | --- | --- | --- |
| BRA | Goat | 1:400 | AF2085 | R&D Systems |
| SOX2 | Rabbit | 1:200 | 5024S | Cell Signaling Technology |
| NANOG | Mouse | 1:400 | 560482 | BD Biosciences |
| SMAD2/3 | Mouse | 1:100 | 610842 | BD Biosciences |
| pSMAD1 | Rabbit | 1:100 | 13820 | Cell Signaling Technology |
| pSMAD2 | Rabbit | 1:200 | 8828S | Cell Signaling Technology |

### Data availability

Scripts used for anaylsis are available from github.com/idse/stemcells. Data are available upon request.

---

**Extended Data Fig 1.**
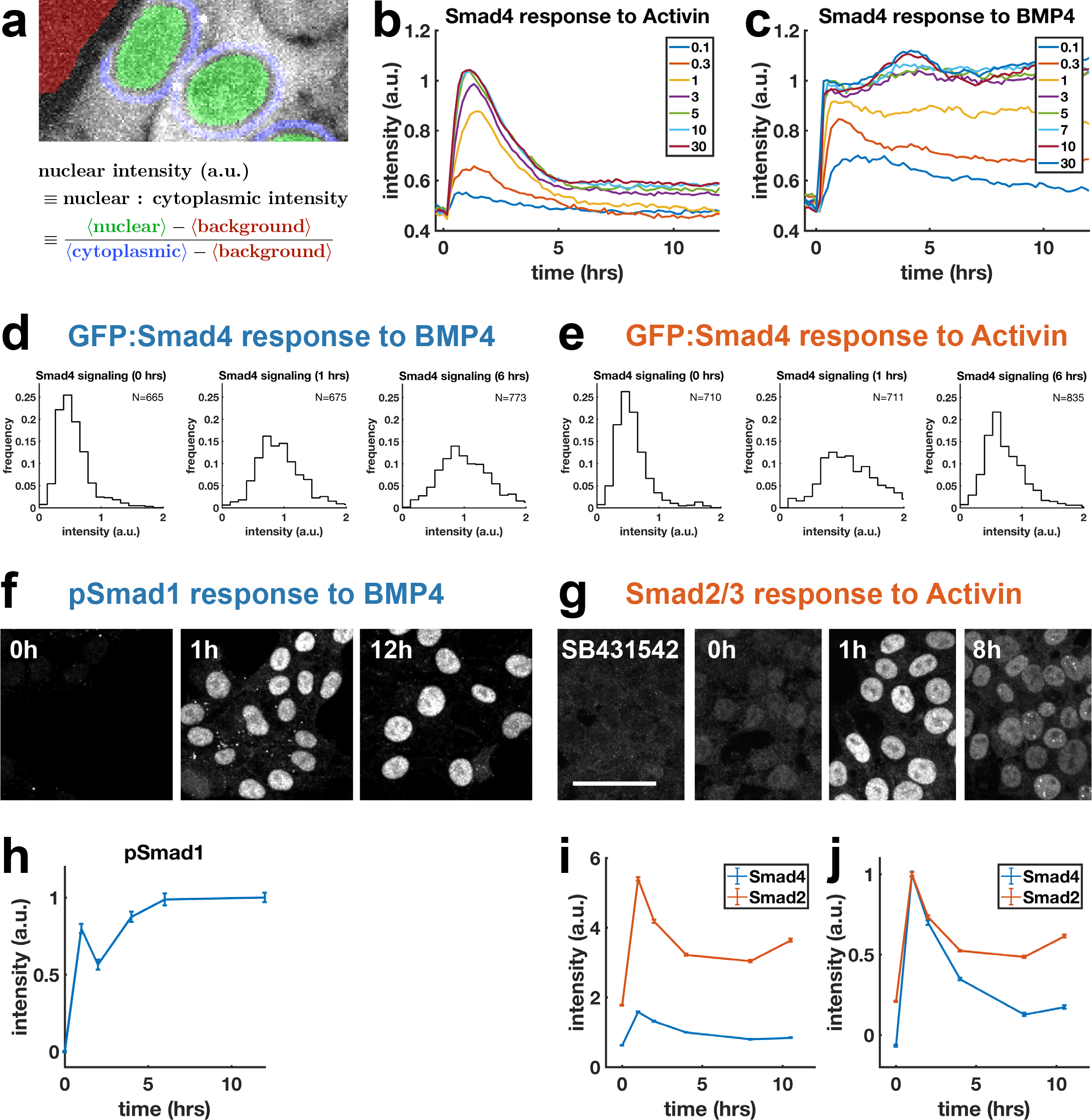
Further characterization of response of hESCs to BMP4 and Activin. **a,** Quantification of SMAD4 signaling as nuclear to cytoplasmic intensity ratio using automated segmentation of the nuclei. **b,** SMAD4 signaling response to different doses of Activin shows fixed time scale of adaptation. Doses in graph legend are in ng/ml. **c,** SMAD4 response to different doses of BMP4 shows decline at low doses with a dose dependent time scale, suggesting ligand depletion. **d,** Distribution of SMAD4 response to BMP4 at 0, 1, and 6 hours after treatment. **e,** Distribution of SMAD4 response to Activin at 0, 1, and 6 hours. **f,** pSMADI time course. **g,** SMAD2 immunofluorescence time course. Scalebar 50 μm. **h,** Quantification of pSmadl time course. **I,** Quantification of SMAD2 and SMAD4 nuclear signaling (nuclear to cytoplasmic ratio). Error bars represent standard error. Ncells per time~10^3^. **j,** Quantification of SMAD2 time course, normalized so that the mean in SB431542 treated cells is zero and peak signaling is 1.

**Extended data fig 2.**
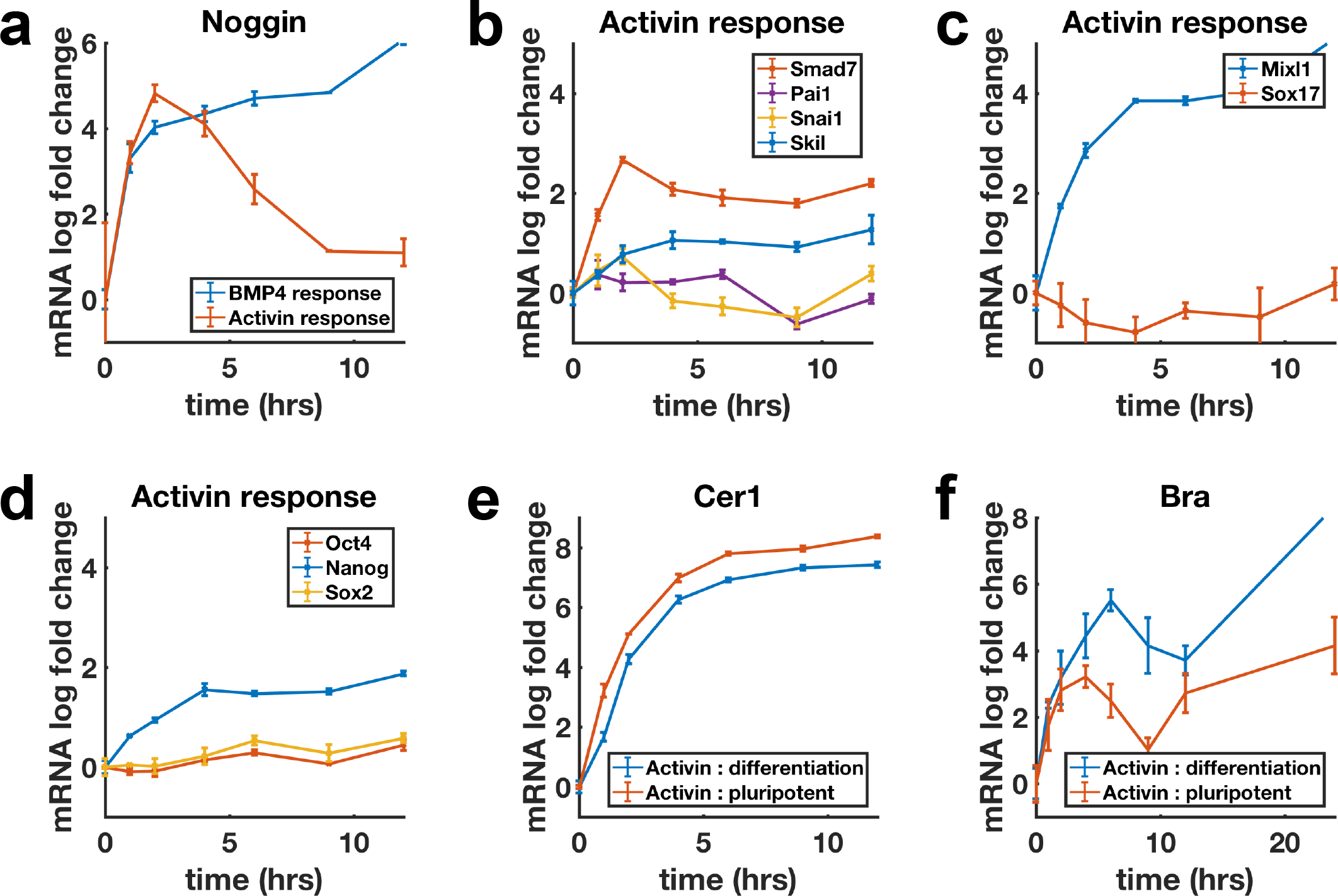
Additional qPCR data. **a,** Transcriptional response of *NOGGIN* to BMP4 (blue) and Activin (red) follows SMAD4 dynamics of respective pathways. **b-d,** Genes in several functional classes show non-adaptive transcriptional response to Activin. **b,** Non-cell fate related (TGF-β targets). **c**, Differentiation genes, *MIXL1* is an exception and responds non-adaptively to Activin, *SOX17* does not respond in the pluripotent state. **d,** Pluripotency genes. **e** *CER1* is a non-adaptive target of Activin that behaves identically under under pluripotency (+FGF) and differentiation (+Wnt) conditions. **f,** Like *EOMES*, *BRA* response is enhanced under differentiation conditions but the dynamics are qualitatively similar.

**Extended Data Fig 3.**
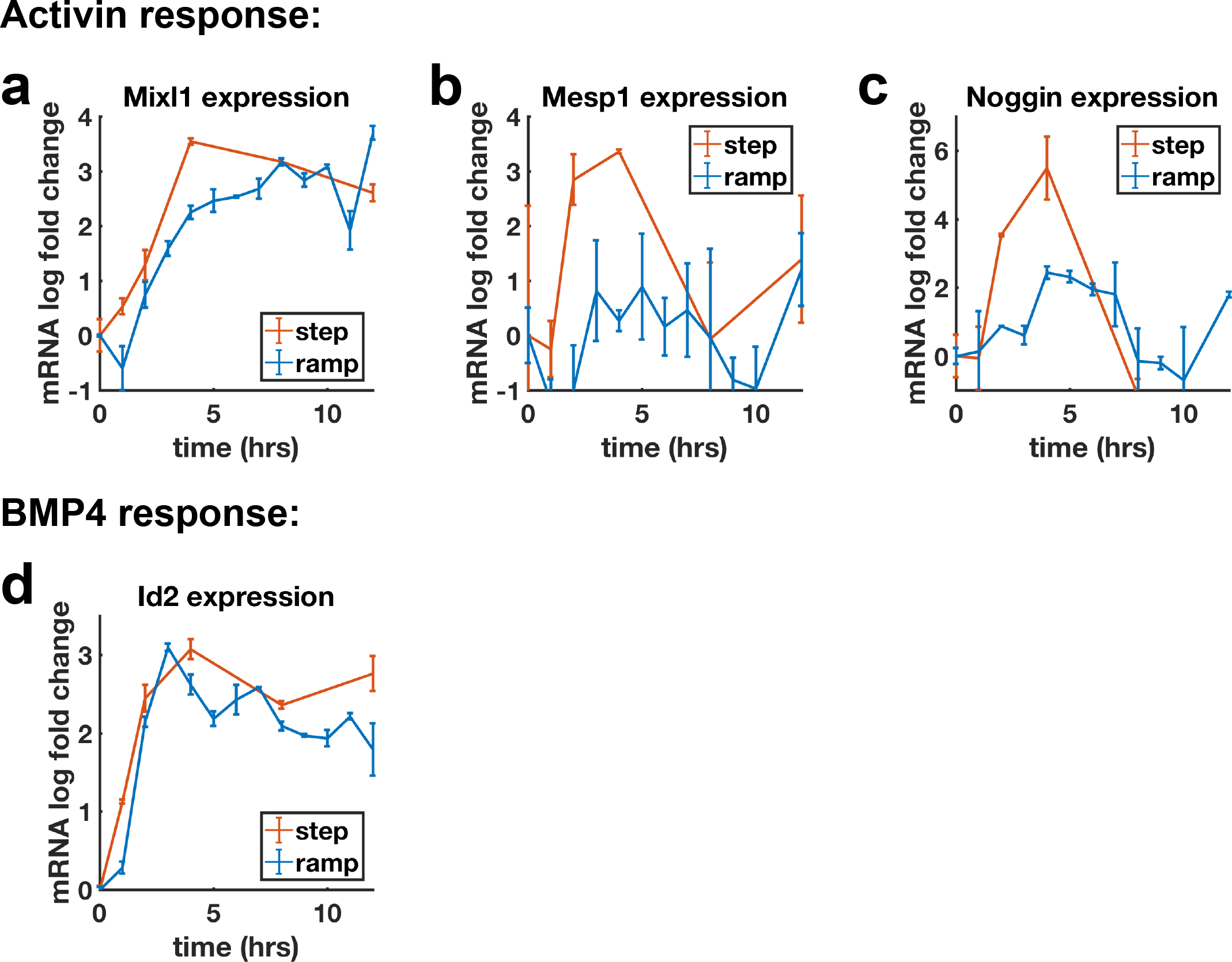
Additional qPCR ramp data. **a,** Transcription of *MIXL1*, a non-adaptive Activin target reaches the same levels after Activin ramp and step. **b,c,** Transcriptional response of *MESP1* **(b)** and *NOG **(c)*** to Activin ramp and step shows reduced transcription for ramp. **d,** Transcriptional response of *ID2* to BMP4 ramp and step.

**Extended Data Fig 4.**
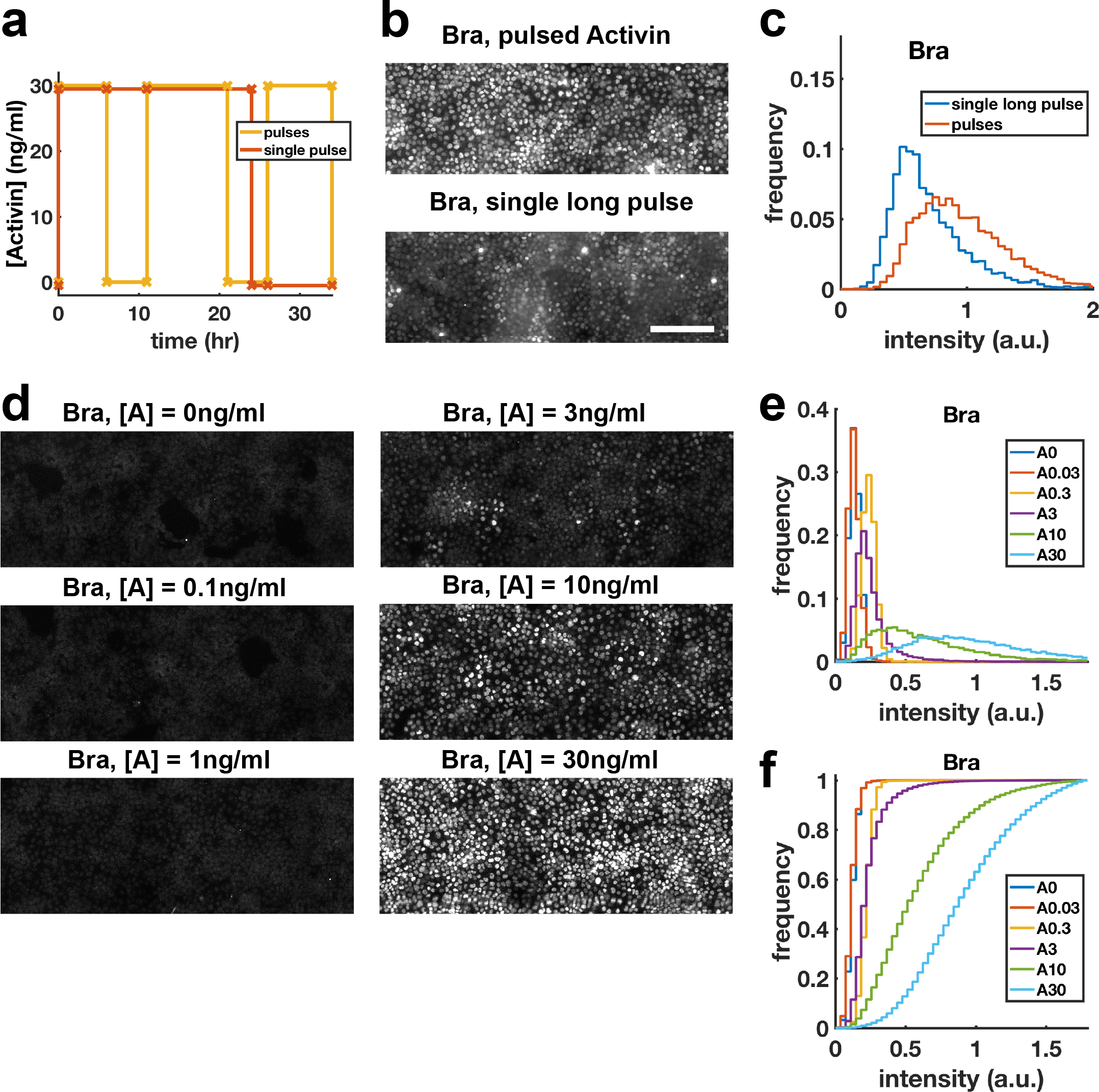
Increased Bra expression after pulsing is not due to reduced integrated exposure. **a-c,** Comparison of BRA expression after single long pulse to 3 shorter pulses with same total time high or no Activin, demonstrating that effect of pulses is not simply due to reduced overall time in Activin. **a,** Schematic of the protocol. **b,** Immunofluorescence stainings for BRA. Scalebar 100μm. **c,** Distribution of BRA intensity per cell in images containing **b** (N_cells_ per condition ~6×10^3^). **d,e,** Dose response series for showing BRA expression monotonically increases with Activin dose, demonstrating that effect of pulses is not due to reduced average Activin exposure. **d,** immunofluorescence staining for BRA after 34h differentiation with different doses of Activin. **e,** Distributions of BRA intensity per cell in the images containing **(d). f.** Cumulative distributions of BRA intensity.

